# Insulin Amyloid Morphology is Encoded in H-bonds and Electrostatics Interactions Ruling Protein Phase Separation

**DOI:** 10.1101/2024.01.10.574993

**Authors:** Samuel Lenton, Hussein Chaaban, Mohammed Khaled, Marco van de Weert, Birgit Strodel, Vito Foderà

**Author notes:** Equal contribution (Samuel Lenton, Hussein Chaaban, Mohammed Khaled).

## Abstract

Ion-protein interactions regulate biological processes and are the basis of key strategies of modulating protein phase diagrams and stability in drug development. Here, we report the mechanisms by which H-bonds and electrostatic interactions in ion-protein systems determine phase separation and amyloid formation. Using microscopy, small-angle X-ray scattering, circular dichroism and atomistic molecular dynamics (MD) simulations, we found that anions specifically interacting with insulin induced phase separation by neutralising the protein charge and forming H-bond bridges between insulin molecules. The same interaction was responsible for an enhanced insulin conformational stability and resistance to oligomerisation. Under aggregation conditions, the anion-protein interaction translated into the activation of a coalescence process, leading to amyloid-like microparticles. This reaction is alternative to conformationally-driven pathways, giving rise to elongated amyloid-like fibrils and occurs in the absence of preferential ion-protein binding. Our findings depict a unifying scenario in which common interactions dictated both phase separation at low temperatures and the occurrence of pronounced heterogeneity in the amyloid morphology at high temperatures, similar to what has previously been reported for protein crystal growth.

## Introduction

Cells utilise membrane-enclosed organelles to selectively segregate specific biochemical reactions within isolated sub-cellular compartments. However, more recently, it has been shown that cells can create transient compartments that can segregate specific components, regulated by a process called liquid-liquid phase separation (LLPS).^1–4^ In protein systems, LLPS is characterised by the formation of two distinct liquid phases, one enriched with the protein, known as the dense phase, and one containing a lower concentration of the protein called the dilute phase.^5^ Due to the highly concentrated protein phase, LLPS provides ideal conditions for the onset of nucleation processes and, in some instances liquid-to-solid transitions. ^6,7^

The nucleation and growth of crystals from protein-dense phases has been reported for a number of proteins. This process has been explained by a two-step crystallisation mechanism, involving the formation of a nucleus that undergoes metastable liquid-liquid phase separation, followed by a conformational change of the proteins that leads to the formation of a highly ordered crystalline phase.^8–11^ This mechanism differs from classical nucleation theory, in which the nucleus is formed directly from a supersaturated solution. ^12,13^ LLPS may influence crystallisation outcomes, such as crystal morphology, and determine the metastable forms of crystals.^14,15^

In addition to crystallisation, and under specific destabilising conditions, the protein-dense phase can undergo a transition into semi-crystalline and gel-like phases;^16^ the latter is generally referred to as aggregation. This transition has been reported for many protein condensates associated with neurological disorders, including FUS in ALS, Tau in Alzheimer’s and *α*-synuclein in Parkinson’s disease. The reaction can result in the formation of amyloid aggregates, ^1–4^ which are characterised by enrichment in beta-sheet secondary structure.This phenomenon also plays a role in drug stability, as certain biopharmaceuticals can also undergo LLPS at the high concentrations required for formulations, resulting in a highly concentrated metastable protein-dense phase. Such metastable phases are generally detrimental to both the stability and efficacy of biopharmaceuticals. ^17^ Furthermore, biopharmaceuticals can undergo amyloid transition, which can result in in vivo complications, such as the formation of insulin balls characterised by large deposits of insulin amyloid at the injection site; ^18,19^ this can result in decreased efficacy and possible immunogenic response to the active compound. These discoveries suggest a new paradigm in our understanding of the origin of protein self-assembly pathways, in which LLPS represents a key step preceding massive aggregate formation.^7^ Indeed, LLPS can affect protein oligomerisation by reducing the translational entropy of proteins^20^ and lead to accelerated kinetics, change in conformational dynamics, and activation of diverse aggregation pathways. ^6^ In the case of intrinsically disordered proteins, the formation of dense protein phases has been shown to act as a precursor for amyloid formation.^21^

Amyloid reactions are regulated by a delicate balance between the hydrophobic attraction caused by protein unfolding and the electrostatic barrier to aggregation, which modulates protein–protein interactions during the onset of amyloid formation.^22^ In this context, the presence of ions in protein solutions can give rise to complex phase diagrams resulting in, for example, coacervation, liquid-liquid phase separation, and reentrant condensation. ^23,24^ When induced by ions, these types of phase behaviours are caused by charge neutralisation or the formation of salt bridges. Although these phase behaviours normally take place below the temperatures used for aggregation studies, their presence sheds light on the interprotein interactions present in solution.^25^ In fact, the magnitude of attractive, short-range interactions between proteins present under dilute conditions has been used to predict the propensity of proteins to phase separate or crystallise at higher protein concentrations. ^26–30^

Ions from the Hofmeister series can solubilise or desolubilise proteins and stabilise or destabilise protein structure. Buell et al. showed that different anions modified protein-protein interactions, thus altering the elongation phase of the aggregation reaction. ^22^ Klement et al. determined that interactions of ions with proteins resulted in modification of surface tension and protein secondary structure, changing the aggregation kinetics. ^31^ Owczarz and Arosio showed that the kosmotropic sulphate anion could delay the aggregation of human insulin at low pH.^32^ In a more recent study, we observed that different anions from the Hofmeister series dictated the final amyloid morphology formed by insulin. ^33^ Moreover, the anion dependency of amyloid morphology could not be described by simple charge screening, suggesting that there are more complex, anion-specific interactions that should be considered.

Here, we provide a unifying framework that connects crystallisation, protein oligomerisation and diversity of amyloid morphologies with the propensity for forming a protein-dense phase, which is controlled by the type and concentration of ions from the Hofmeister series. Using microscopy, small-angle X-ray scattering and atomistic molecular dynamics simulations, we demonstrate that, in solution, the sulphate and perchlorate anions induce phase separation of insulin by specific binding to the protein. Sulphate ions also determine the oligomerisation state of insulin and, upon storage at room temperature, the crystal formation. Upon incubation at high temperatures, the ion specific binding favours a coalescence process over a purely conformation-driven aggregation, leading to the occurrence of morphologies other than elongated fibrillar structures. As in crystallisation, a protein system’s propensity to undergo LLPS dictates the type of nucleation mechanisms and aggregate morphology under amyloid-forming conditions.

## Results

### Anions from the Hofmeister series regulate the cloud point temperature of human insulin

The cloud point temperature of human insulin at pH 2.0 was assessed by titrating the sodium salt of each anion directly into a known concentration of insulin. During titration, the sample temperature was fixed and both the absorbance in the UV region and the light scattering at 90° were recorded. When the cloud point temperature was reached, the insulin solution became visually turbid and this turbidity was reflected in a sudden increase in both the measured scattered intensity and the UV absorbance at 600 nm. Control measurements were conducted for buffer solutions without insulin, and no increase in scattering was detected throughout the examined temperature range. Calculating the final concentration of both insulin and titrated anion at this point yielded the phase diagrams shown in Fig. 1a and b. Regarding the chloride anion, no sudden changes in turbidity were observed for the concentrations of anion and insulin tested. However, for both sulphate and perchlorate, the cloud point temperature is dependent on both the anion and insulin concentration (Fig. 1a, b). Whilst investigating the insulin solutions, we also observed that solutions of certain concentrations became turbid at low temperatures (4 °C) but returned transparent after heating them back to room temperature. Therefore, we also performed titration experiments at 10 °C, whereupon we observed a shift in the phase boundaries to lower concentrations (Fig. 1a, b).

**Figure 1:**
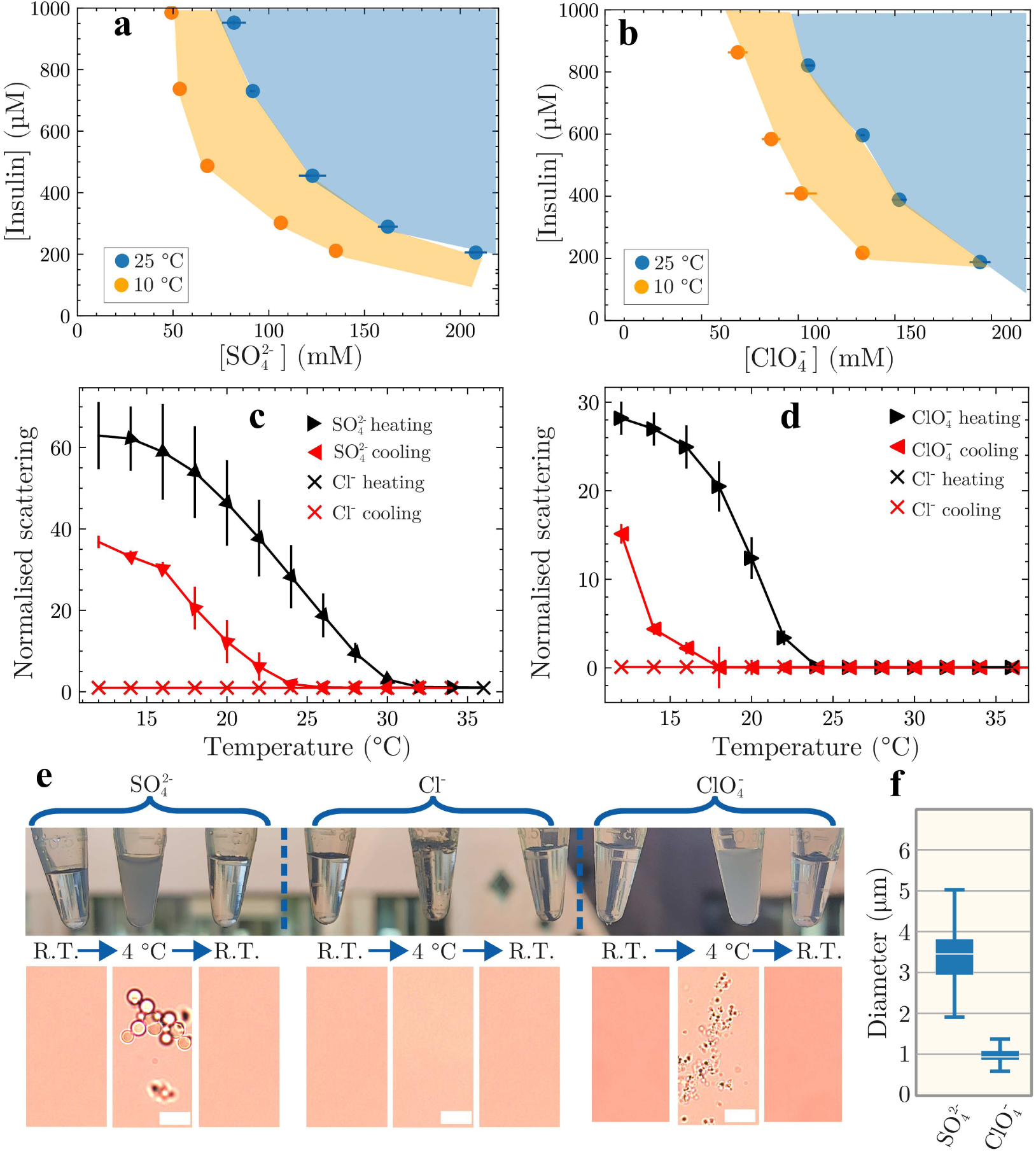
**(a, b)** Insulin phase diagrams as a function of insulin and anion concentrations at the indicated temperatures shown for **(a)** 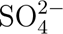 and **(b)** 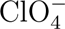, determined by measuring scattered light intensity at a fixed angle. Shading indicates the region above the cloud point temperature for the respective concentrations. **(c and d)** Turbidity curves of insulin (600 µM) in the presence of **(c)** 100 mM 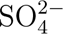 and **(d)** 100 mM 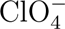 shown as a function of temperature and measured by light scattering at a fixed angle. **(e)** Top row: Photographs of insulin solutions before cooling, when cooled to 4 °C and upon heating them back to room temperature. Bottom row: Bright-field images of the solutions under the respective conditions (scale bar=10 µm) **(f)** Size distribution of the formed coacervates (*n* at least 50 coacervates measured for both samples).

### Both perchlorate and sulphate induce temperature-dependent phase separation of insulin

To investigate the reversible nature of the temperature-induced turbidity, we measured the scattering intensity at 90° as a function of temperature (with 2 °C steps and 3 minutes of equilibration for each step) for both the sulphate and perchlorate anion at a fixed insulin concentration (Fig. 1c and d). For both samples, an increase in scattering was observed with decreasing temperature. This was accompanied by an increase in sample turbidity, which decreased upon increasing the temperature and finally returned to a transparent state. In presence of sulphate, the solutions became completely transparent at 30 °C, while for perchlorate, this happened at around 25 °C. The same solutions were then cooled again and an increase in scattering was observed at 22 °C and 17 °C for sulphate and perchlorate, respectively. It should be noted that after sufficient incubation time at 8 °C (15 additional minutes), both samples returned to their original turbidity indicating complete reversibility (not shown).

The same experiment performed with 100 mM chloride showed no increase in turbidity at any of the measured temperatures (cross symbols in Figs. 1c and d). This result indicates an upper critical solution temperature (UCST)-type behaviour for insulin in the presence of 100 mM sulphate and perchlorate but not chloride.^34^ The observed hysteresis between the heating and cooling curves suggests multiple interactions of the sulphate and perchlorate anions with insulin, resulting in changes of both inter- and intra-protein interactions.^34^

To investigate the causes of the observed turbidity, a single insulin sample was prepared for each anion and divided into three Eppendorf tubes. For each anion condition, one tube was kept at room temperature, while the other two were cooled to 4° C for 30 minutes; of these, one was heated back to room temperature for 15–20 minutes (Fig. 1e, Fig. S1,2). The samples were then observed using light microscopy. As expected, for the sample containing sodium chloride, no significant change with temperature was observed, whereas the turbid sulphate and perchlorate samples contained droplets (Figure 1e). We verified that the observed structures were protein-enriched by centrifuging the samples, which resulted in two distinct phases: a transparent top phase and a turbid bottom phase. For both samples, UV measurements of the top phase indicated that they were depleted of protein and they did not show any features under the light microscope. There was a significant difference in the size of the droplets formed, in the order of 3.4 *µ*m and 1 *µ*m for sulphate and perchlorate, respectively (Figure 1f). Interestingly, in sulphate-containing insulin solutions, the growth of protein crystals was observed at room temperature after 48 hours (Fig. S3).

### Anion-type and concentration determine the oligomerisation state

Having established the bulk properties of the insulin solution, and having identified anion-specific UCST-type behaviour, we wanted to gain a deeper understanding of the effects that the selected anions may have on the oligomerisation properties of insulin. To study this in the solution state, small-angle X-ray scattering was measured on freshly prepared insulin solutions (i.e., at time 0). Measurements were performed at room temperature for a range of insulin and anion concentrations. We note that room temperature lies above the UCST of both the sulphate and perchlorate solutions, thus, for these measurements, the protein is homogeneously diluted in the solutions. Model-independent analysis was performed by plotting the data in the Kratky depiction and determining the radius of gyration from the Guinier plot. Representative Kratky plots for the chloride anion (100 mM) at two insulin concentrations are shown in Fig. 2a. Under the same salt conditions, increasing the insulin concentration resulted in a shift of the maximum towards lower *q*. Due to the reciprocal nature of the SAXS data, this indicates the formation of larger particles in the solution.

**Figure 2:**
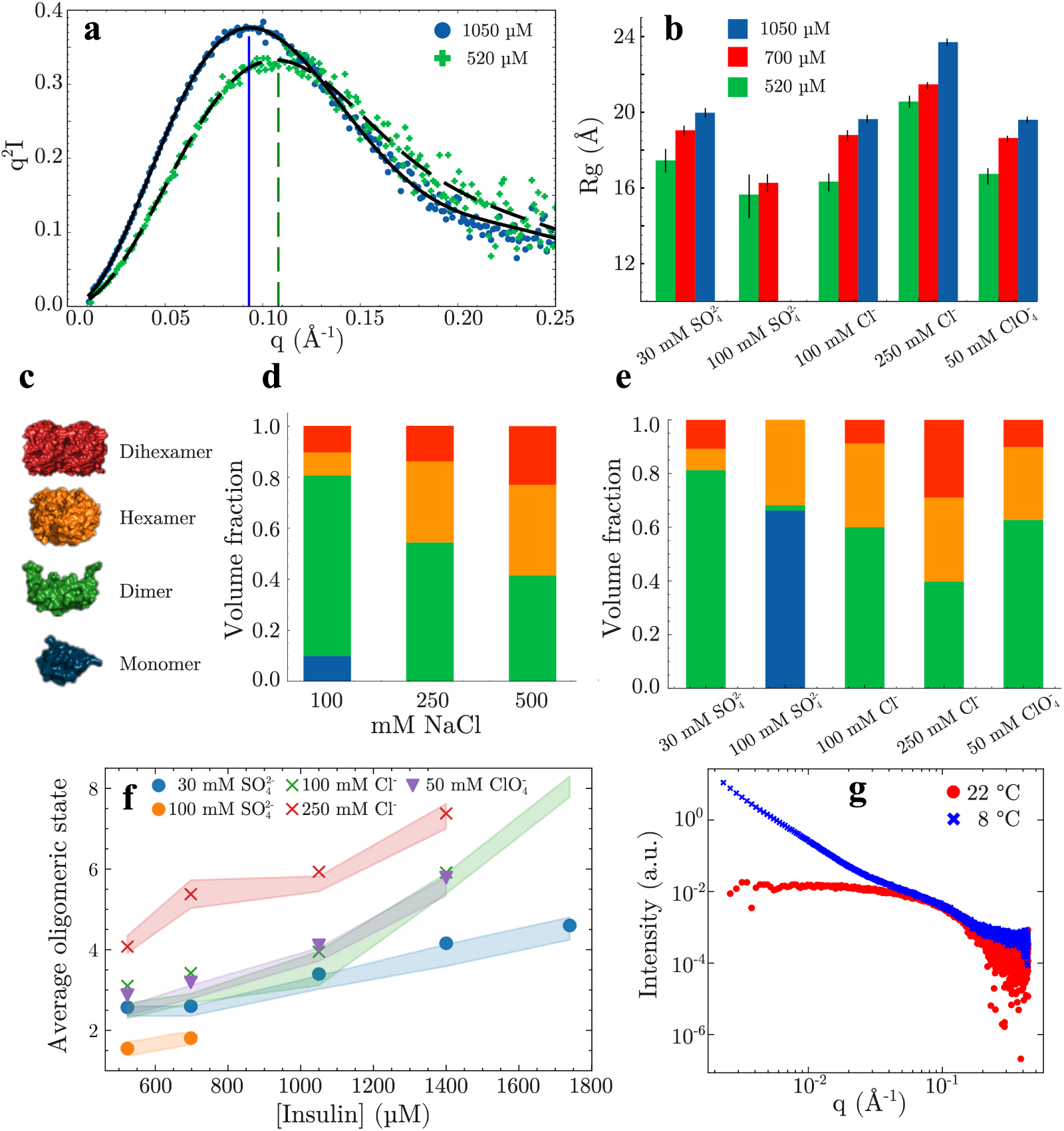
**(a)** Example Kratky plot of the insulin SAXS data at the indicated insulin concentration with 100 mM NaCl. The dashed line indicates the maxima, which shift to lower *q* with increasing insulin concentration. The concentration of insulin is indicated in the legend. **(b)** Radius of the gyration of insulin samples under different ionic conditions at the indicated insulin concentration, determined from the Guinier fit to the SAXS data. **(c)** Depiction of insulin structures fitted to the SAXS data. **(d)** Distribution of oligomers determined from fitting of the SAXS data for 520 *µ*M insulin at the indicated NaCl concentrations. The colours correspond to the oligomeric species shown in (c). **(e)** Distribution of oligomers determined from fitting the 4 mg/ml SAXS data shown to the indicated anion solutions; colours correspond to the oligomers shown in (c). **(f)** The average oligomeric state of insulin determined from the SAXS data based on the calculation of the oligomeric distribution (symbols) and the calculation of the volume directly from the SAXS data (shaded areas). **(g)** SAXS spectra of insulin (520 *µ*M) before cooling (red) and after cooling at 8 °C for 15 minutes (blue).

This insulin concentration-dependent increase in the size of the protein particles in solution was confirmed by an increase in the radius of gyration (*R*_g_), which was determined from a linear fit to the Guinier region of the SAXS data; the gyration radii obtained are shown for all anions at two insulin concentrations in Fig. 2b. Although an increase in *R*_g_ with increasing insulin concentration was observed for all anion conditions, both the *R*_g_ value at the lowest insulin concentration and the magnitude of the observed increase were anion-dependent.

Insulin tends to form a range of oligomeric species (e.g., dimers, hexamers, Fig. 2c) in solution.^35^ Thus, we questioned whether the changes in radius of gyration were due to the extent of oligomerisation. To test this hypothesis, we calculated theoretical scattering curves of different insulin oligomers from crystal structures and fitted them to SAXS data.^36^ Good fits to the scattering data were obtained for most conditions, indicating that well-defined insulin oligomers were present in solution. For the chloride anion, SAXS experiments and analysis were performed at three different anion concentrations and at a fixed insulin concentration of 520 *µ*M (Fig. 2d and Fig. S4). These results indicate that larger oligomeric species formed with increasing ionic strength from 100 to 500 mM. This is in agreement with the expected increase in protein-protein interactions upon increasing the screening of charges in solution. At the highest protein and anion concentrations measured, the fits obtained started to deviate from the experimental data because of the presence of oligomers or aggregates larger than those used to fit the data.

The average molecular mass was calculated from the theoretically obtained distribution of oligomeric species (Figs. 2e and 2f). To further verify that the oligomeric distribution obtained was accurate, the molecular mass was also calculated based on the volume calculated from the SAXS data (i.e., without fitting the insulin oligomeric structures) and compared with the molecular mass determined from the calculated oligomeric distribution (Fig. 2f). In general, a strong agreement was found between the two methods, supporting our conclusion that insulin oligomers were formed. Comparison at a single insulin concentration with varying anions (Fig. 2e) revealed that the chaotropic anion perchlorate resulted in similar distributions of oligomeric species compared to the intermediate Hofmeister-series chloride anion. However, the kosmotropic sulphate anion showed distinct effects. At 100 mM sulphate, a large fraction of monomeric insulin was present, which was not observed for any of the other anionic species or concentrations (Fig. 2e). We next investigated the effect of the critical temperature on the insulin solutions containing sulphate. The scattering curve obtained at room temperature depicts the one expected for insulin (Fig 2.g). The sample was next cooled below the critical temperature (8 °C for 15 minutes), and SAXS was measured again. The resulting SAXS spectra shows a massive increase in the measured scattering intensity in the low *q* range, indicating the formation of the large clusters observed in Fig. 1e.

Based on observations from the the SAXS analysis, we conclude that, for all anions, there was insulin concentration-dependent oligomerisation, with more insulin participating in larger oligomers with increasing concentration. For the neutral chloride anion, a clear influence by the ionic strength on the extent of oligomerisation was observed, as expected of a purely electrostatic interaction. However, sulphate was an outlier, as a decrease in average molecular mass was observed with increasing sulphate concentration. We further observed that at 100 mM sulphate, oligomerisation was inhibited and most of the insulin in solution was in monomeric form.

### Anions change the thermal stability of, and interact with, insulin in an anion-specific manner

To understand the specific ion effects of the selected anions and their interactions with insulin in more detail, we performed atomistic molecular dynamics (MD) simulations at varying concentrations of 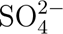, Cl*^−^* and 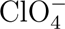. Simulations were conducted at 25 °C and 65 °C, with the higher temperature selected to resemble the insulin solution properties under the aggregation conditions reported previously. ^33^ The stability of proteins hinges on the stability of the secondary structures within the encompassing tertiary structure. We therefore determined the secondary structure propensity of each residue throughout the MD simulation according to temperature (25 or 65° C). These propensities were then subtracted from each other to yield the so-named delta secondary structure percentage (Figs. 3a, b and c), which depicts the change in secondary structure elements upon heating. For both chloride and perchlorate anion, a significant decrease in the *α*-helical nature of the first ten N-terminal residues was observed, while for sulphate, there was an increase in *α*-helical structure in this region. Between the C-terminal region of chain A and the N-terminal region of chain B, there was an increase in beta sheet-type structure in the presence of chloride. For perchlorate, there was no significant change in this region, although, for sulphate, there was a slight loss in random coil and an almost equal increase in *α*-helix. The changes in the secondary structure upon heating can be summarised as follows: perchlorate demonstrated a loss of *α*-helical secondary structure in the N-terminal region and the same effect was observed for chloride coinciding with an increase in beta sheet. For sulphate, there was a slight increase in the *α*-helical structure in the N-terminal of both chains. For both perchlorate and sulphate, there was no detectable appearance of beta sheet upon heating.

**Figure 3:**
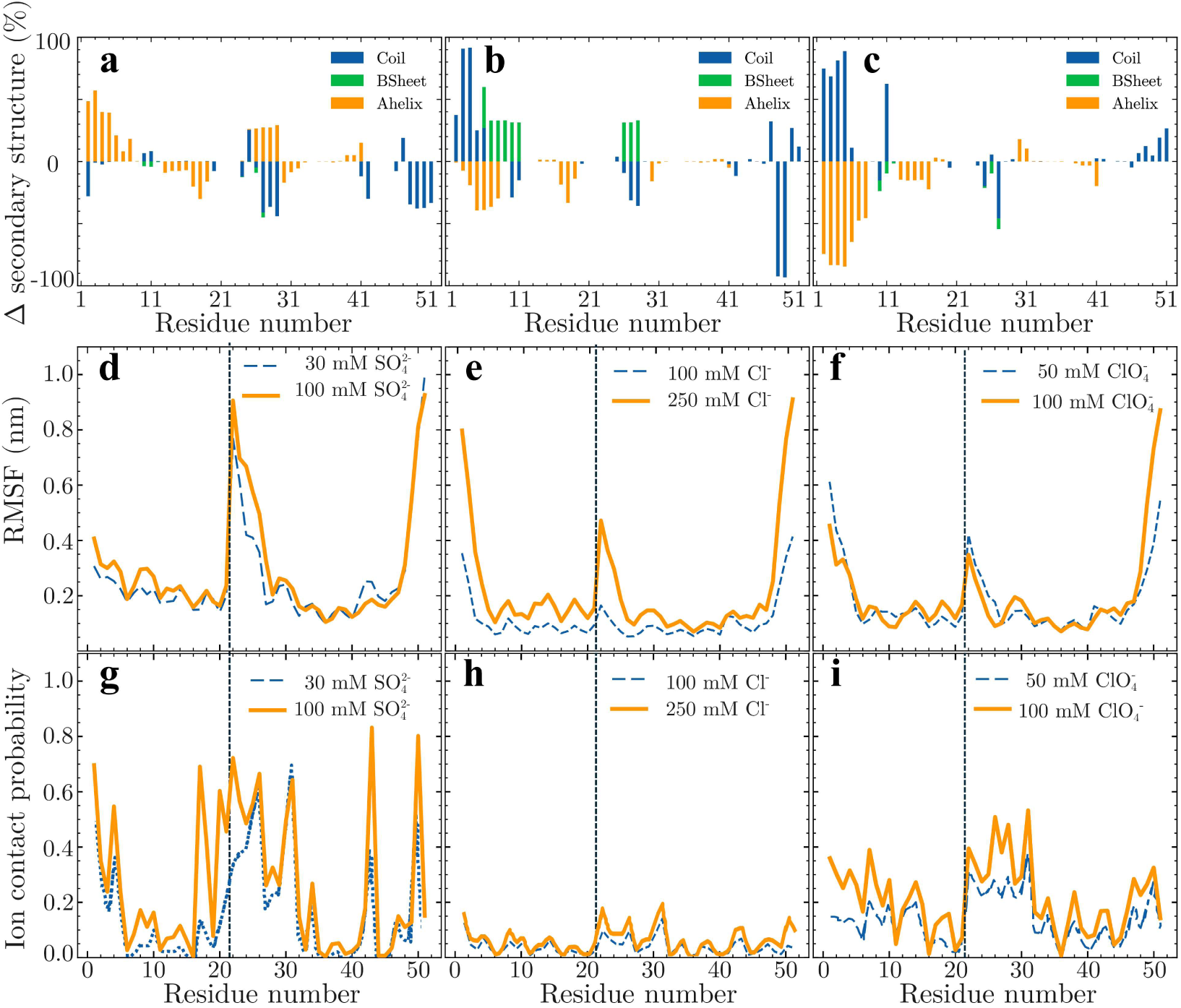
Results from the insulin MD simulations in the presence of the selected anions. **(a, b, c)** Change in insulin secondary structure observed upon heating to 65° C for (a) 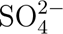, (b) Cl*^−^*(c) 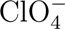. **(d,e,f)** The root mean square fluctuations (RMSF) of C*α* carbon atoms, at the ionic condition indicated in the legends, from the 25° C simulation. The dashed vertical line indicates the terminus of the a-chain. **(g, h, i)** Ion contact probability for each residue with anion and concentration indicated, from the 25° C simulations. The dashed vertical line indicates the terminus of the a-chain.

The root-mean-squared fluctuations (RMSF) determined from the MD simulations are shown in Figs. 3d, e and f. At the start of chain B (i.e., residues 22-30), there was an increase in the RMSF for all three conditions. The fact that this region fluctuated more than the rest of the insulin protein is not surprising considering that this region is known to be flexible.^37^ More significantly, this region fluctuated more heavily in the presence of the sulphate anion compared to in the presence of perchlorate and chloride anions. Increased fluctuations were also observed at the N-termini of chain A and the C-terminus of chain B, corresponding to unstructured flexible regions in the crystal structure of insulin (Figs. 3d, e and f).

To determine the extent to which specific ion-protein interactions differ between the selected Hofmeister anions, the ion contact probability on a per-residue basis was extracted from the MD simulations. For all anion conditions, there was no significant probability of ion contact between any of the insulin residues and the cation (Na^+^). Similarly, the chloride anion did not show significant tendencies to interact with the insulin protein (Fig. 3h). On the contrary, for both perchlorate and sulphate, an increase in the probability of the ion interacting with insulin was observed at the C-terminal of chain A and at the N-terminal of chain B (Figs. 3g and i). This high anion contact probability, the highest for sulphate followed by perchlorate, and the lowest for chloride, coincided with a prolonged ion lifetime on the same residues with a high contact probability (Figs. S5, S6 and S7). Comparing the three anions, we can conclude that there was a ranking of anion interactions with the protein, in order of increasing interaction strength, of: chloride, perchlorate, and sulphate.

### Hydrogen bonding underpins the strong anion-insulin interaction

From the MD simulations, the average number of hydrogen bonds per insulin residue at 25 and 65° C was extracted (Fig. 4a, b and c). The results show that sulphate formed several hydrogen bonds with insulin, while both perchlorate and chloride showed negligible hydrogen bond formation with insulin. On the basis of the apparent propensity for hydrogen bond formation between insulin and sulphate, we hypothesised that a combination of hydrogen bond formation and electrostatic interactions causes the tight interaction between sulphate and insulin. Compared with perchlorate, this tight interaction is likely increased because of the divalent negative charge and the presence of multiple hydrogen bond acceptors (the oxygens), allowing sulphate to form bidentate hydrogen bonds to adjacent proteins (Fig. 4d). The formation of at least two hydrogen bonds between the sulphate and adjacent insulin monomers could act as a bridging interaction that results in the formation of a protein network. If present, this interaction could explain the enhanced propensity of sulphate to form droplets at a lower temperature, and explain the differences observed in the droplet size, between perchlorate and sulphate (Figure 1f).

**Figure 4:**
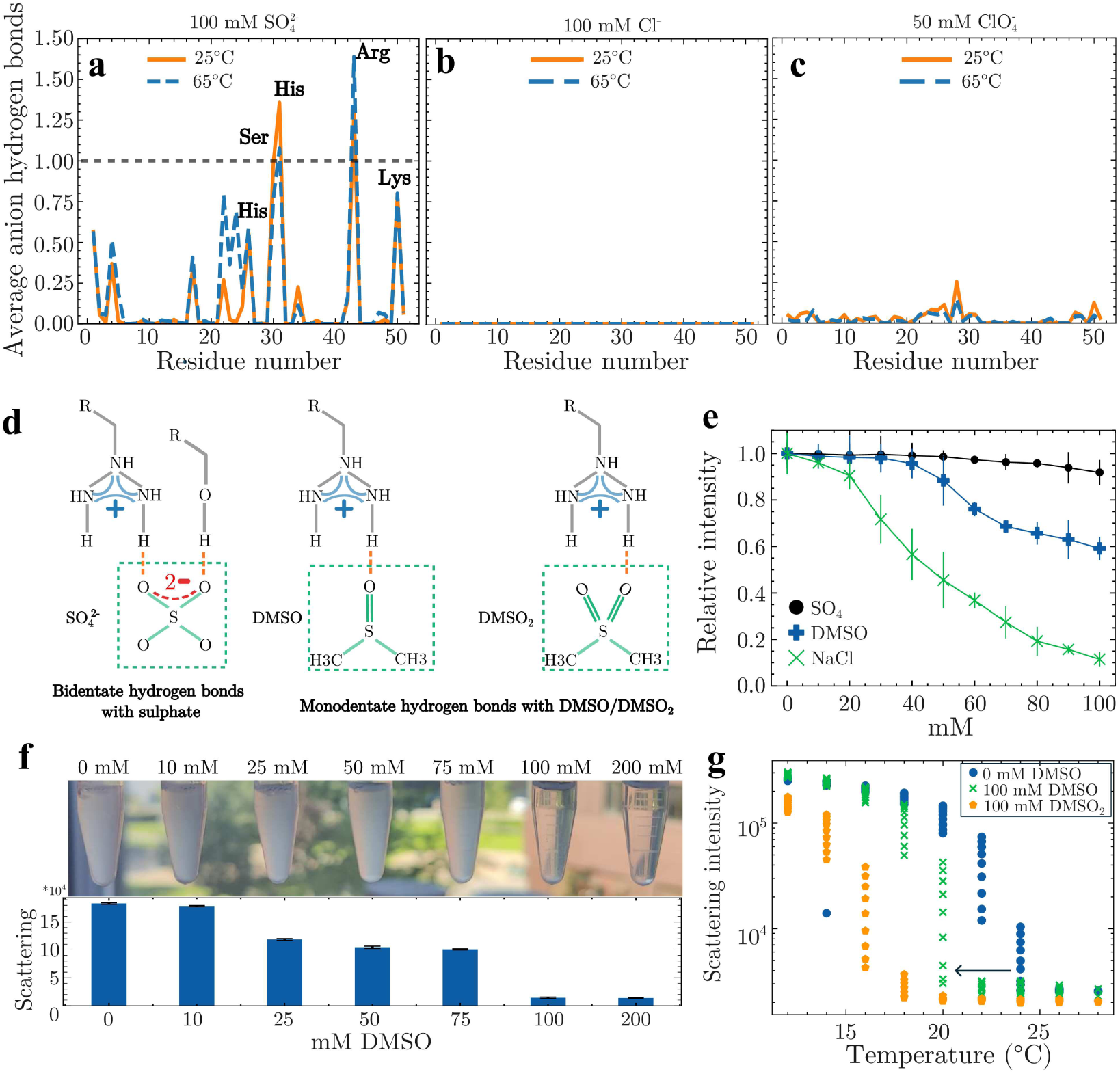
**(a, b and c)** Average number of hydrogen bonds between the residues of insulin and the indicated anions, extracted from the MD simulations. **(d)** Schematic depiction of potential hydrogen bond pairs between amino acid side chains and the indicated excipients **(e)** Effect of 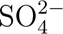, DMSO and NaCl concentration on the turbidity of an insulin sample (600 *µ*M) in 100 mM 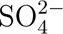 in the phase-separated state. **(f)** Top: Pictures of insulin solutions (600 *µ*M) in 100 mM 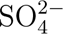, with increasing concentration of DMSO. The solutions were incubated at 8° C for 15 minutes. Bottom: Light scattering intensity measured for the corresponding samples shown in the top row. **(g)** Cloud point measurements performed for 600 *µ*M insulin solutions containing 100 mM 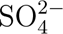, in addition to the indicated concentration of excipients. Measurements were performed by decreasing the temperature.

To test the contribution of both electrostatic and hydrogen-bond interactions to the observed droplet formation, we titrated sodium chloride and Dimethylsulfoxide (DMSO) into an insulin solution below the critical solution temperature (i.e., when dense droplets occur) whilst measuring the turbidity by scattering intensity. DMSO has been shown to act as a strong hydrogen bond-accepting competitor that disrupts the polymer networks formed by hydrogen bonds (Fig. 4d).^38^ The addition of sodium chloride, on the other hand, increases the total ionic strength of the solution, which increases the Debye length and strengthens the screening of electrostatic interactions. We note that the DMSO concentrations used here are far below the concentrations typically required for the unfolding of human insulin, so that the changes reported below are not due to unfolding processes.^38^ Since the titration experiments involves a dilution of the solution, control experiments were also performed with an equivalent concentration of sulphate solution to account for any potential effects in the experiment solely due to dilution.

The results of the titration experiment are shown in Fig. 4e. Upon titration of sulphate into the turbid insulin solution, only a small decrease of intensity was observed, indicating that turbidity persisted upon slight dilution during the experiment. Titration of both DMSO and sodium chloride resulted in a dramatic decrease in the measured turbidity, with sodium chloride having a stronger effect than DMSO. These data indicate that it is a combination of electrostatics and hydrogen bonding that contributes to the interactions that stabilise the observed droplets. Having tested how DMSO influences the droplets of already formed solutions, we next prepared fresh insulin solutions at a fixed concentration (600 *µ*M) above the critical temperature, containing increasing concentrations of DMSO. These solutions were then cooled below the critical temperature for 15 minutes. As expected, a decrease in turbidity was observed with increasing DMSO concentration (Fig. 4f), with 100 mM DMSO producing completely clear solutions. This result contrasts with the data shown in Fig. 4e, where turbidity persisted at 100 mM DMSO. This can be explained by the two different experimental protocols used. Indeed, in Fig. 4e, DMSO was added to solutions already containing droplets (i.e., ion–insulin H-bonds already formed) while, in Fig. 4f, DMSO was added prior to hydrogen bond formation and formation of the dense phase.

In a final experiment, we wanted to monitor the impact of H-bonds on the cloud point of insulin, so we performed the experiment on insulin solutions in the presence of 1) 100 mM sulphate, 2) 100 mM sulphate and 100 mM DMSO, and 3) 100 mM sulphate and 100 mM DMSO_2_. Compared to DMSO, DMSO_2_ contains an additional oxygen available for H-bond formation (Fig. 4d). Although cloud point temperatures of 24 and 20 °C were detected for sulphate and DMSO, respectively, in the presence of DMSO_2_, this value further decreased to 16° C, demonstrating a decisive influence of the ion-protein H-bonds on the formation of the insulin-dense phase.

In summary, the interaction between sulphate and insulin is caused by a combination of electrostatic interactions and hydrogen bonding. Reducing the contribution of these interactions resulted in a decrease in the droplet phase. The formation of the droplet phase is caused by the ability of both perchlorate and sulphate to form bidentate hydrogen bonds, resulting in the crosslinking of a large number of insulin monomers. However, compared to perchlorate, sulphate solutions exhibited a higher critical temperature and formed larger droplets. This can be explained by the divalent charge of sulphate, which enhances the hydrogen bond formation with the protein, resulting in tighter anion binding and the formation of larger protein networks.

### Anions determine aggregation kinetics and amyloid morphology

The aggregation kinetics of insulin at 65 °C in the presence of the three Hofmeister anions are shown in Fig. 5a. Compared to both perchlorate and chloride, sulphate displayed a slower growth rate that coincided with a prolonged period of time to reach the plateau phase. The overall kinetic profile observed for sulphate differed significantly from the sigmoidal pathway observed for classical amyloid fibrils. ^39,40^ The prolonged delay time and reduced kinetics were consistent with previous experimental data, showing that, with sulphate, large aggregates without beta-sheets were initially formed, which converted over time to ThT-positive beta-sheet aggregates. ^32^ The formation of microparticles, is evident in the microscopy images (Fig. 5b.) taken at the end of the aggregation process, showing ThT-positive, sphere-like structures without any birefringence, and is in agreement with our previous work.^33^ Sodium chloride resulted in sigmoidal aggregation kinetics with a lag phase followed by rapid growth, resulting in the formation of ThT-positive amyloid fibrils (Fig. 5c.). Compared to chloride, perchlorate caused a slight increase in the initial lag phase, followed by a rapid growth phase, resulting in the formation of predominantly spherulite structures, characterised by the presence of a Maltese-cross pattern under polarised light, with some amyloid fibrils (Figs. 5d and e.). This co-existence is in agreement with previous studies. ^41^

**Figure 5:**
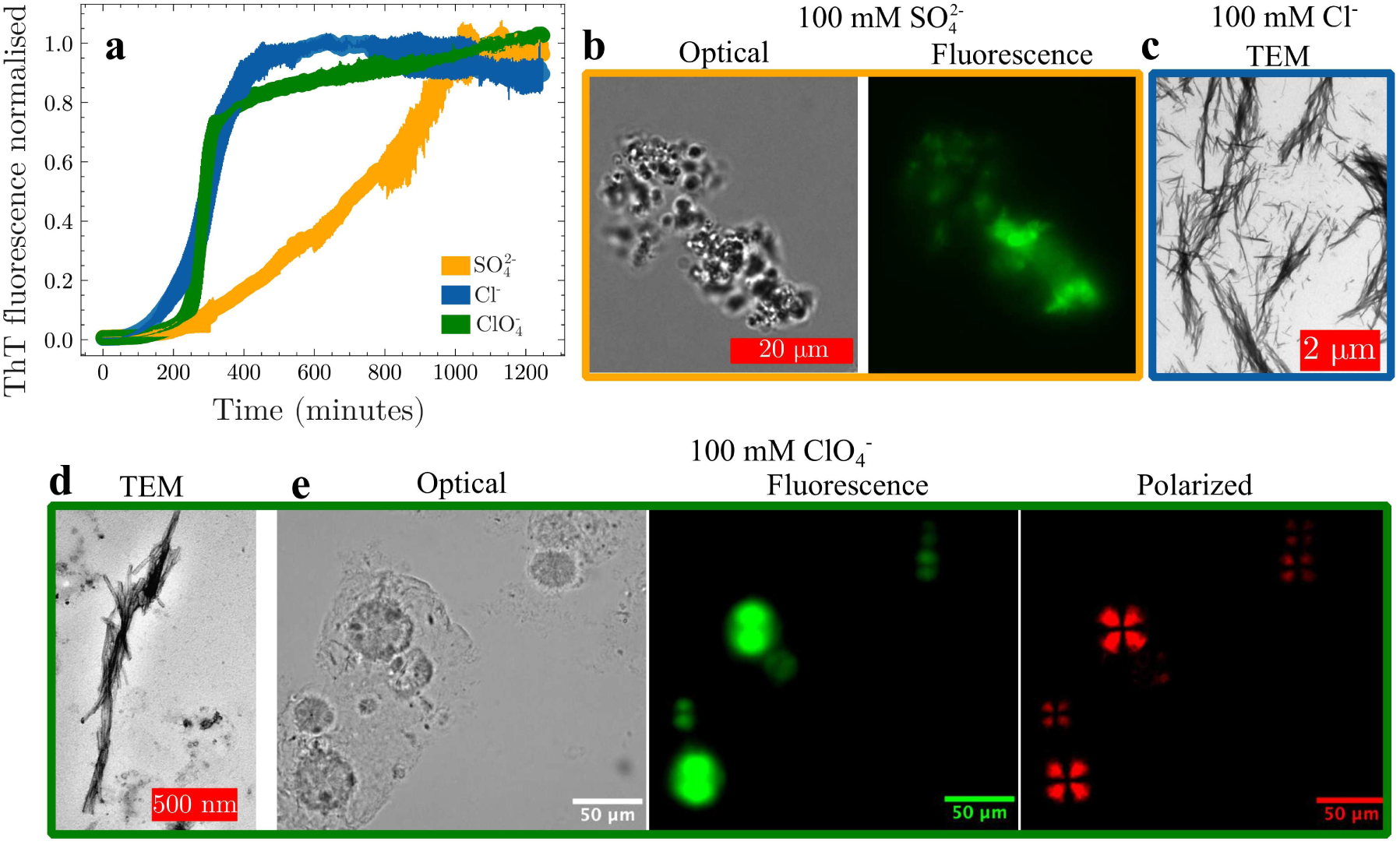
**(a)** Aggregation kinetics of 600 *µ*M insulin at pH 2 and 65 °C in the presence of 100 mM of the indicated anion, measured by ThT fluorescence. **(b)** Amyloid particulates formed in 100 mM 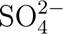 and stained by Thioflavin T, as observed through optical and fluorescence microscopy. **(c)** TEM of amyloid fibrils observed with 100 mM Cl*^−^*. **(d)** TEM of amyloid fibrils observed with 100 mM 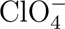. **(e)** Optical microscopy, ThT fluorescence, and polarised light microscopy of insulin with 100 mM 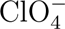, stained with Thioflavin T. The polarised light microscopy images reveal a Maltese cross pattern characteristic of spherulitic amyloid aggregates.

## Discussion

Proteins may self-assemble into various forms of amyloids, from elongated fibrillar structures and dense spheres, called particulates, to hedgehog morphologies called spherulites, resembling a structure commonly encountered in polymer science. ^33,39,42^ These structures have been observed both in vivo and in vitro, and are considered general pathways for a number of proteins, independent of their amino acid sequence. ^43–45^ Electrostatic interactions between proteins are the major determinant of this heterogeneity of amyloid structures, but the precise mechanism that leads to this diversity is still unknown. In this work, we used a selection of Hofmeister anions below concentrations where one would expect to see the kos-motropic and chaotropic nature of Hofmeister anions (typically above 300 mM). Despite the low concentrations used, the selected Hofmeister anions regulated the formation of distinct amyloid morphologies (Fig. 5).^33^

To determine how these different anions exert such a drastic influence on the observed aggregation phenomena, we first studied their effects on the solution properties of insulin prior to aggregation. We observed that, in the presence of both sulphate and perchlorate, but not chloride, insulin solutions displayed an upper critical solution temperature, below which formed droplets are enriched with protein (Fig. 1). Compared to perchlorate, sulphate had a higher critical solution temperature and, at the same temperature, formed larger protein droplets and thus had a stronger effect on the solution properties of insulin.

At the molecular level, the data from the MD simulations showed that heating both the perchlorate- and chloride-containing systems caused a decrease in the *α*-helix content of insulin, and the presence of chloride even induced the formation of beta sheets (Fig. 3). On the other hand, in the case of sulphate, there was only a slight increase of *α*-helix upon heating and an absence of beta sheet formation. This observation corroborates previous experimental data, which showed that sulphate can lock insulin in the monomer state with an enhanced conformational stability against temperature increases. ^32^ Further examination of the MD data showed that the interactions between the anions and insulin occurred in order of increasing interaction strength: chloride, perchlorate and sulphate.

For the monovalent and monoatomic anion chloride, no specific ion contact probability was observed with insulin and, unsurprisingly, no specific hydrogen bonds formed (Fig. 3). For perchlorate an increase in the ion contact probability with insulin was predominantly found at the first six N-terminal residues of the B-Chain (Fig. 6). In the case of sulphate, a notably increased probability of contact was detected, which was associated with an extended duration of ion interaction and a significant level of hydrogen bond formation. These interactions were mainly located in the N-terminal region of the B-chain, as well as in the area involved in dimer formation of the B-chain, and the N-terminal segment of the A-chain (Fig. 6). We confirmed that this enhanced capacity of sulphate binding to insulin was due to a combination of electrostatics and hydrogen bond formation using titration with both DMSO and NaCl (Fig. 4). This also explains why UCST behavior was also observed with perchlorate, but was not as pronounced as with sulphate. This results in the formation of smaller droplets by perchlorate, as sulphate is both divalently charged and can act as a multiple hydrogen bond acceptor. Previously, the crystal structure of insulin has been determined under similar solution conditions (i.e. pH 1.7 in the presence of sulphate). ^46^ After incubating the sulphate-containing insulin solutions for 48 hours at room temperature, we also observed the growth of protein crystals (Fig. 7 and Fig. S3). The insulin structure determined from the sulphate-containing crystals by Whittingham et al. showed the presence of several sulphate molecules bound to specific residues of insulin. Several of these residues are consistent with the data from our MD study (Fig. 6). In the crystal structure, the sulphate ions also bridge neighbouring insulin monomers in the crystal, highlighting the ability of sulphate to act as a cross-linker between adjacent proteins.

**Figure 6:**
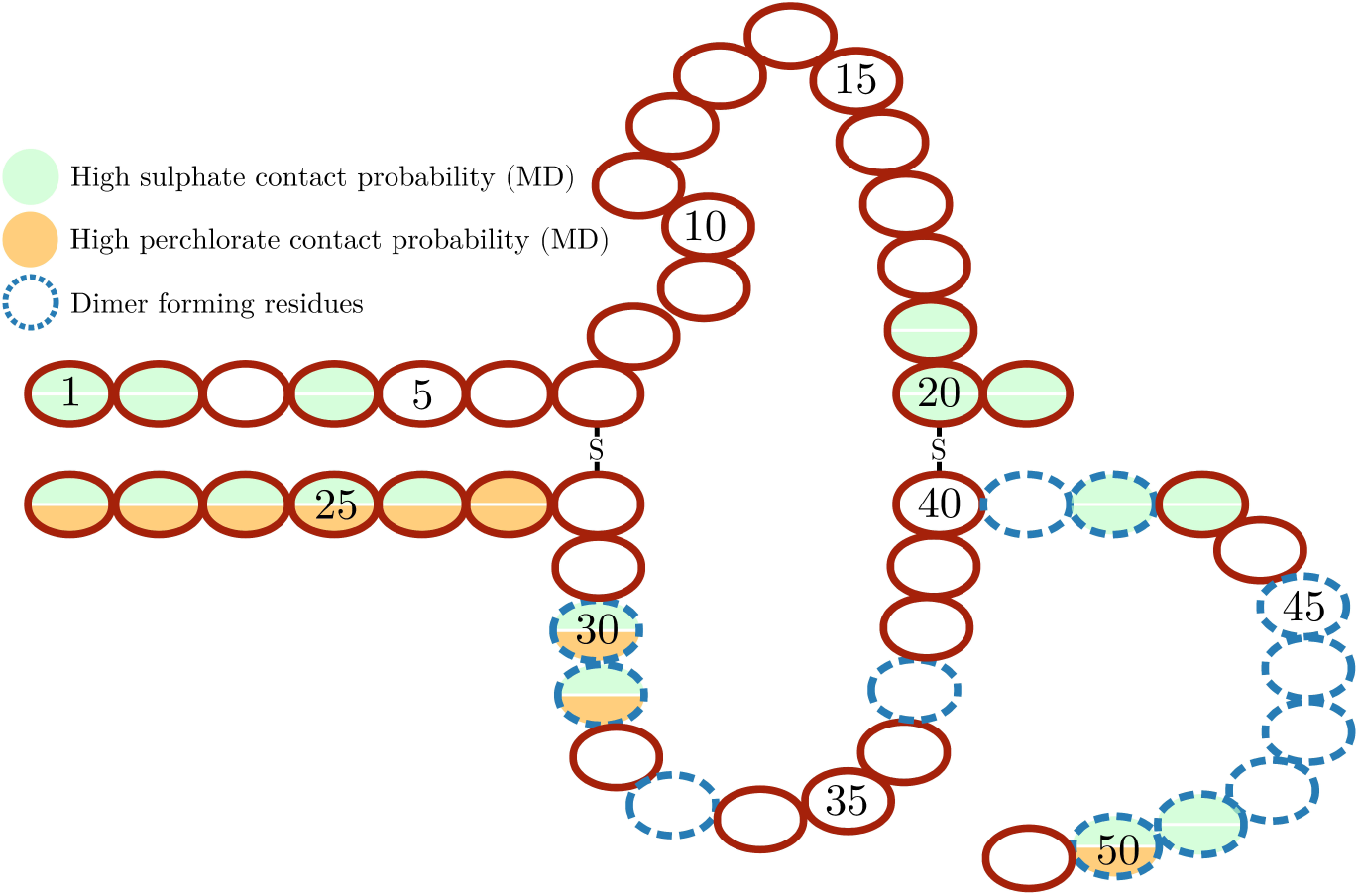
The primary structure of insulin depicting the A- and B-chain linked together by disulphide bonds. Residues with an ion contact probability determined by the MD simulation to be more then 0.4 are highlighted for perchlorate and sulphate.

**Figure 7:**
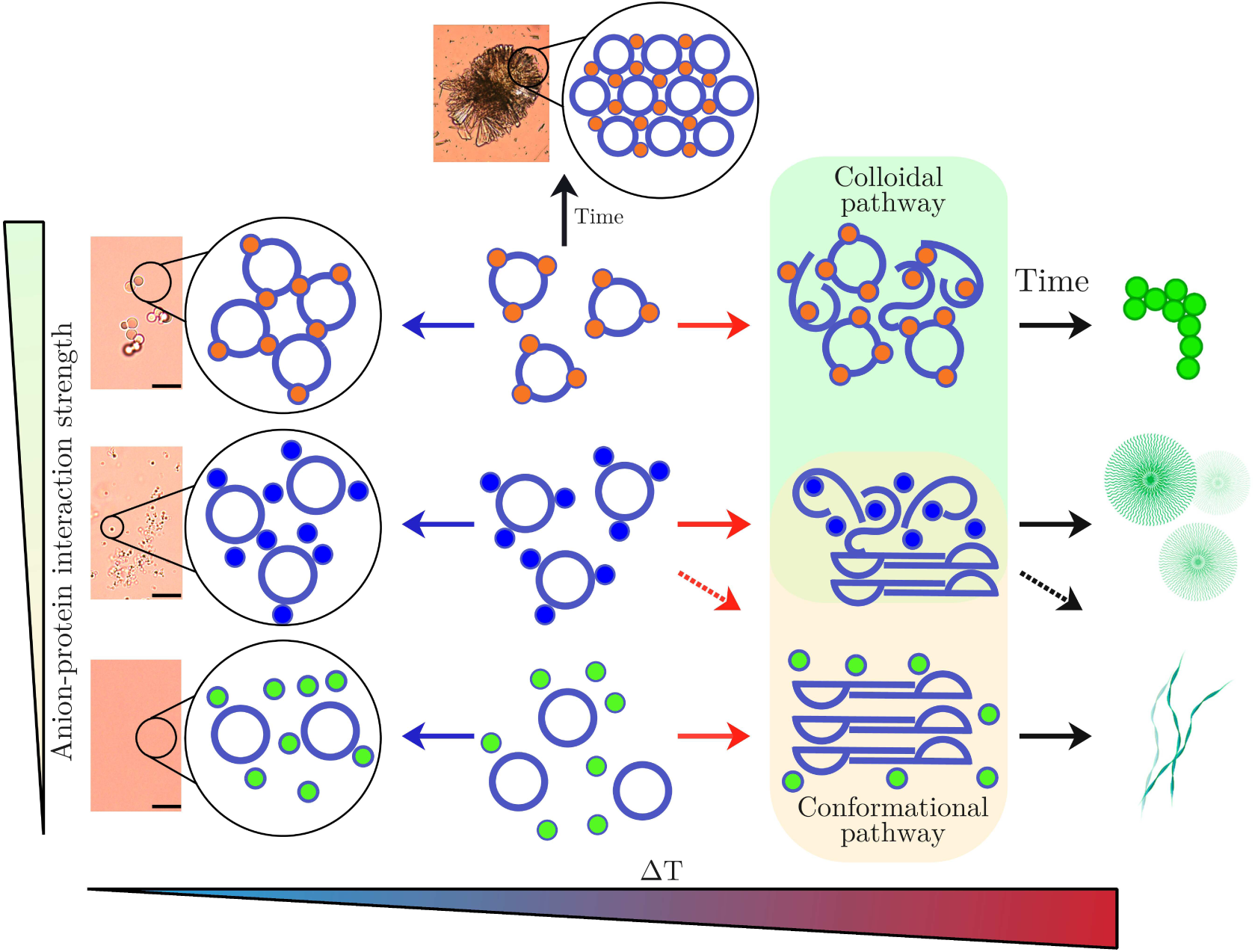
Schematic overview of the results. **Top row:** Colloidal pathway imposed by sulphate ions that interact specifically with insulin resulting in a reduction of the net positive surface charge. Upon heating, larger aggregates are formed that retain the folded structure. Over time, these aggregates convert to the beta-rich structures observed in amyloids. Attractive inter-protein interactions can also be observed when the mixture is cooled, resulting in phase separation, and when left at room temperature, causing the formation of protein crystals. **Middle row** Perchlorate results in an intermediate between the colloidal and conformational pathways. There is a reduced UCST behaviour, and no protein crystals are observed after incubation at room temperature. Aggregates form but grow predominantly through the recruitment of unfolded monomers, resulting in the formation of predominantly spherulite structures. **Bottom row** Conformational pathway imposed by chloride anions. The chloride anions do not interact specifically with the protein and instead provide charge screening based on ionic strength. The formation of amyloid is driven by the unfolding of the insulin protein, resulting in the formation of beta sheet-rich elongated fibrils.

After determining that hydrogen bonding of sulphate, strengthened by electrostatic interactions, affects the phase behavior and crystallization of insulin, we proceeded to examine the solution properties of insulin at room temperature at the initial time point. We observed both anion-specific and concentration-dependent changes in the average oligomerisation state of insulin (Fig. 2). In general, larger oligomers were formed with increasing concentration of protein and anion. However, for sulphate, we observed a distinct lack of oligomers even at higher protein concentrations, with the predominant species being the insulin monomer. We propose that this effect may also be caused by the tight binding of sulphate, specifically to the dimer-forming region which prevents further oligomer formation (Fig. 6). In the case of the crystal formation, it was suggested that this specific binding of sulphate to insulin acts to neutralise the surface charge of the protein, thus propagating the crystallisation process. Previous experimental studies of lysozyme have observed a similar effect. Gokarn et al. measured the charge of lysozyme at relatively low ionic concentrations, such as below 100 mM.^47^ They observed a decrease in the effective charge of lysozyme following the order: chloride, perchlorate, sulphate. They attributed this decrease in effective surface charge to increases in specific binding of sulphate compared to the other anions.

In order to understand how the specific interactions of the anions influence the final morphology of amyloids formed, we performed a range of microscopy experiments. Based on the results in Fig. 5 and previous experimental data, the influence of the anions on the final amyloid morphology is as follows: sulphate results in mainly amyloid-like spherical microparticles (particulates), chloride produces mainly elongated amyloid fibrils, and perchlorate produces a mixture of fibrils and spherulites.^33^ At low pH insulin has an overall net positive charge, resulting in a long-range electrostatic repulsion between monomers. However, if specific anion binding occurs at the surface of the protein, the overall net charge is decreased, reducing the long-distance electrostatic repulsion between insulin molecules allowing them to approach each other. At low temperatures, the insulin molecules then coalesce and form the observed droplets. Generally, for globular proteins, raising the temperature tends to accelerate amyloid formation because of the necessary unfolding step. Importantly, in the case of sulphate, ion binding enhances the conformational stability of the protein (Fig. 3). In this configuration, it should be expected that the aggregation process will exhibit features similar to a coalescence of colloidal particles, with the protein initially maintaining its structural arrangement and approaching neighbouring protein molecules. Indeed, our ThT fluorescence data (Fig. 5) show that the formation of structures rich in beta sheet proceeded more slowly in the presence of sulphate compared to either perchlorate or chloride. This result is in line with the work of Owczars and Arosio, who showed that, under conditions similar to those studied here, the lag time of insulin aggregation approximately doubled when the anion was changed from chloride to sulphate (i.e., 2.5 vs 5 hours).^32^ In the same study, solutions of insulin and sulphate showed a dramatic increase in hydrodynamic radius prior to the onset of beta sheet formation. This means that, already in the early phase of the process, larger aggregates are formed, which retained their *α*-helical structure for a prolonged period before undergoing a beta-sheet transition. Interestingly, the link between the degree of conformational stability and the diversity of aggregation pathways was also shown for the protein concanavalin A.^48^

This process can be explained by the presence of a locked monomeric state of insulin and reduced electrostatic repulsion, induced by the tight binding of sulphate to the surface. Together, these two aspects result in a colloidal-like aggregation reaction in which structurally unchanged proteins coalesce and form spherical particles. This pathway, termed the colloidal pathway, is depicted schematically in Fig. 7. The microparticles formed by coalescence eventually undergo beta-sheet transition at later stages, as previously observed for both insulin and *α*-lactalbumin.^32,39^ We propose that the tight binding of sulphate to insulin, due to a combination of hydrogen bonding and electrostatic interactions and resulting in a reduced net charge and an enhanced conformational stability, is at the basis of both the appearance of the protein-dense phase at low temperatures and the formation of amyloid particulates at high temperatures.

Under aggregation conditions in the presence of chloride anions, we mainly observed the formation of amyloid fibrils. Previous work reported a full correlation between scattering and ThT kinetics, suggesting that the massive aggregation observed in chloride solutions is paralleled with an *α*-helix-to-beta sheet transition.^49^ We term this traditional pathway the conformational pathway (Fig. 7).

Finally, in the case of perchlorate, the situation lies somewhere between the other two. Compared to chloride, perchlorate binds more tightly to insulin, with slight hydrogen bond formation, but not as strongly as sulphate, resulting in a weaker UCST-type behaviour (Fig. 1). Upon aggregation of insulin in perchlorate solutions, a more complex scenario emerges, and both amyloid fibrils and spherulites are observed. The latter are described as a core-shell structure, characterised by a radially growing beta-sheet aggregates on a disordered precursor core. ^50,51^ This apparent heterogeneity of amyloid morphologies can be explained by an interplay between a purely colloidal pathway and a conformation-driven reaction. Indeed, at higher temperatures, the electrostatic repulsion between insulin monomers is still reduced, but not to the same extent as for sulphate anions. This fact allows for the formation of smaller aggregates by coalescence. However, the conformational stability of the insulin-perchlorate system is also reduced compared to the sulphate sample (Fig. 3), allowing a higher propensity towards conformational changes. At high temperatures, this last aspect has two main consequences. In one case, unfolded proteins undergo a purely conformation-driven reaction, similar to the case of the insulin–chloride aggregation and resulting in the formation of amyloid fibrils. As a second pathway, the aggregates formed by coalescence may act as a nucleation site for unfolded proteins, which, in turn, promotes the formation of radially growing aggregates, eventually defining the fingerprint of spherulites (Fig. 7).

In summary, we demonstrate that the specific interactions of anions from the Hofmeister series regulate not only the phase behaviour and stability of insulin but also the solid transitions to crystalline and amyloid forms. We report that the interactions responsible for the droplet formation of insulin at low temperatures are the determinants of the solid transition at higher temperatures, both in terms of crystals (at room temperature) and amyloid aggregates at higher temperatures. In the latter case, the interactions regulating the dense phase are also responsible for the balance between colloidal and conformational stability of the protein, leading to diverse pathways and a pronounced heterogeneity in the structure and morphology of the amyloid aggregates.

In the context of neurodegenerative diseases, the aggregate polymorphism is associated with different patterns of neuropathology,^52,53^ while the adverse biological effects of protein drugs depend on the structure of the aggregate. ^54–56^ Our findings address the concept of morphological fingerprint of aggregates encoded in the LLPS propensity of a protein. This perspective may guide a more tailored screening of modulators for protein aggregation, especially if specific amyloid polymorphs are targeted. The identification of protein condensed phases will indeed serve as a rapid prediction method for potential aggregate structures and conformations. In addition, short-lived intermediate species and oligomers are indicated as targets for pharmacological treatments, but their isolation is often challenging, even in in vitro experiments.^57^ In this regard, protein-dense phases will provide a more accessible and long-lived protein ensembles for testing molecules that can inhibit or reverse the aggregation process, paving the way for novel opportunities in the treatment of degenerative disorders and protein drug development.

## Methods

### Chemicals

Human insulin (91077C, 95%), sulphate (99%), sodium chloride (99%), sodium perchlorate (98%) were purchased from Sigma-Aldrich, Germany. Sodium hydroxide (99%) and hydrochloric acid (99%) were purchased from Merck, Germany Th.Geyer and CHEMSOLUTE, Denmark, respectively.

### Cloud point temperature measurements

Fresh samples of insulin were prepared by dissolving a known amount of insulin and setting the pH to 2.0 by titration of HCl. The samples were then filtered through a 0.22 *µ*m syringe filter (Labsolute, Th.Geyer, Germany) and the final concentration of insulin was determined by triplicate measurements of U.V. absorbance with a Nanodrop system (Thermo Fisher). The freshly prepared samples of insulin were then placed in a quartz cuvette with a stirring bar and incubated at the specified temperature in a Labbot instrument (ProbationLabs, Sweden). The autotitration system of the instrument was used to slowly titrate 1 M of either sulphate, perchlorate or chloride into the insulin sample, whilst simultaneously measuring the scattering at 90 degrees and the U.V. absorbance in the range 280-600 nm. The data were plotted as a function of salt concentration, after correcting for the dilution effect due to the titration. The concentrations at which a jump in the scattering at 90 °(or the absorbance at high wavelengths) took place was determined to be the cloud point temperature. Each measurement was repeated at least 3 times.

### Temperature dependence measurements – single-angle light scattering experiments

Fresh insulin samples were prepared as before at pH 2, in 100 mM sodium sulphate, perchlorate or chloride. After filtering, the samples were placed in a quartz cuvette with a stirring bar in the Labbot instrument and allowed to equilibrate for at least 30 minutes at 12 degrees. During the experiment the stirring was turned on and the temperature ramped by 2 degrees every 3 minutes. During the temperature run the scattering intensity of a 636 nm laser at 90°angle was measured. The scattering was then normalised by the lowest measured scattering intensity for each sample.

### Titration measurements – single-angle light scattering experiments

Fresh insulin samples were prepared as before at pH 2, in 100 mM sulphate. After filtering, the samples were placed in a quartz cuvette with a stirring bar in the Labbot instrument and allowed to equilibrate for at least 30 minutes at 12°C. During the experiment the stirring was turned on. Every 5 minutes ten microlitres of 1M NaCl, 1M DMSO or 1M Na_2_SO_4_ was titrated into the solution, after which the light scattering intensity was measured at 90°.

### Microscopy measurements

For the detection of the protein rich droplets, samples were prepared by dissolving the insulin in the desired anion concentration and setting the pH to 2.0. After preparation the samples were filtered through a syringe filter (0.22 *µ*m) and the insulin concentration was confirmed by triplicate measurements of the U.V. absorbance at 280 nm. The samples were then left to equilibrate at 4°C for at least 30 minutes before imaging. One of the samples was, however, first heated back to room temperature for a period of about 30 minutes before imaging. 15-30 *µ*l of the samples were then placed on a glass slide and covered with a 22×22 mm cover slip and directly imaged with a 64x oil objective (Zeiss, Germany) using a Leica DMi8 optical microscope (Leica Microsystem, Wetzlar, Germany) using either brightfield, fluorescence (excitation 488 nm and emission at 530 nm) or polarized light settings. The morphology of the protein aggregates (spherulites and particulates) and crystals was detected by using a DMi8 optical microscope (light microscope, LM) with a 10x, 20x and a 63x oil objective (Leica Microsystem, Germany). 10-15 *µ*l of the aggregates/crystals were placed on a glass slide prior to imaging and analyzed in polarized, brightfield and fluorescence modes.

### Data Collection Small-angle X-ray scattering (SAXS)

5 mg of HI powder was dissolved in 1 mL of 30 and 100 mM sodium sulphate, 100; 250 and 500 mM sodium chloride; and 50 and 100 mM perchlorate. Sodium hydroxide and hydrochloric acid were used to adjust the pH to 2.00 ± 0.05. Afterwards, the samples were filtered through a 0.22 *µ*m cellulose filter (Labsolute, Th.Geyer, Germany) and the final concentration of insulin determined by U.V. absorption at 280 nm. 20-30 *µ*l of each sample was transferred to a 96-well plate. The SAXS studies were performed at the CPHSAXS Facility, University of Copenhagen, using a BioXolver L (Xenocs) with a wavelength of *λ* = 1.34 Å. The SAXS experiment with cooling was performed at the P12 beamline operated by EMBL Hamburg at the PETRA III storage ring (DESY, Hamburg, Germany). At DESY, the ARI-NAX BioSAXS sample changer was used to automatically load the samples, while the cold sample was maintained at 8 degrees within the sample holder before being loaded. Primary data reduction was made in BIOXTAS RAW.^58^ Experiments were conducted at room temperature, with samples prepared immediately before the experiment and then stored on ice until loading the sample. Scattering from the corresponding buffers was measured under identical conditions (i.e. same exposure time) and subtracted from the sample scattering.

### Data analysis – SAXS

For each solution condition the radius of gyration was determined by linear fitting to the Guinier region (Ln(I) against *q*^2^) that satisfied *q* · *Rg <*= 1.3. The SAXS data were fitted using the program OLIGOMER from the ATSAS package^59^, using the crystal structures of insulin and insulin oligomers taken from the PDB (PDB ID: 3i40,1guj,1ai0). ^46,60,61^ From the distribution of oligomers the average molecular mass was calculated. The Mw was further calculated from each scattering curve using the DatBayes method implemented in PrimusQT from the ATSAS package.^59,62^

### Aggregation kinetics monitored by ThT fluorescence

A stock solution of 1 mM ThT was prepared by dissolving ThT in milli-Q water and filtering through a 22 *µ*m syringe filter (Labsolute, Th.Geyer, Germany). ThT was added to the solutions containing insulin at 600 *µ*M and a 100 mM NaCl/Na_2_SO_4_/NaClO_4_, to a final concentration of 20 *µ*M. 200 *µ*L of each protein solution was transferred to a 96-well plate (NUNC, Thermo Fisher, USA). ThT fluorescence was measured on a CLARIOstar (BMG Labtech, Germany) plate reader using 450 nm for excitation and a 490 nm filter for detection using the bottom optics. At least 4 repetitions were made for each sample.

### Transmission Electron Microscopy (TEM)

The morphology of the fibrils was observed by TEM using a Philips CM 100 TEM (Philips, The Netherlands) operated at voltage of 80 kV. Images were recorded with the ITEM software package. 5 *µ*l of the sample were negatively stained and mounted on hydrophilic grids (by glow discharge) prior to imaging. For details on sample preparation, please refer to previous work.^63^

### Molecular dynamics (MD) simulations

The initial structure of HI was obtained from PDBs 2mvc.^64^ To mimic the experimental pH 2 level conditions, protonated Lys, Glu, and His were used for modelling the insulin. Also, charged N- and C-termini were used (NH3^+^, COO*^−^*). The insulin carried an overall charge of +5. The HI was placed in a cubic box of 400, 280 nm^3^, with 1.5 nm from the box walls. Each system was solvated by TIP3P water models, then different Hofmeister salt (sodium chloride, sodium sulphate and sodium perchlorate) concentrations were added to mimic the experimental conditions and neutralize the systems. The All-atom MDs were performed using GROMACS 2020 simulation package^65,66^ with Amber14SB forcefield^67^ and the parameters of the ion developed by Kashefolgheta and Baaden for sulphate and perchlorate, respectively.^68,69^ The systems were first energy minimized using the steepest descent minimization algorithm to remove any local atomic clashes. ^70,71^ Subsequently, the systems were equilibrated under an isothermal–isobaric NPT ensemble for 100 ps, where the desired temperatures (25 or 65°C) were achieved using a velocity-rescale thermostat method with a coupling time constant of 0.1 ps.^72^ While the pressure was kept at 1 Pa using the Parrinello-Rahman pressure coupling method with a coupling time constant of 1.0 ps.^73–75^ During the equilibration process, a position restraint with a force constant of 1000 kJ/mol.nm^2^ for protein heavy atoms was applied. The production runs followed, which were performed under NPT ensemble using Nose-Hoover^76–79^ thermostat and a Parrinello–Rahman barostat with a relaxation time of 2 ps. The particle mesh Ewald method (PME) was used to compute the electrostatic interactions.^80,81^ The cutoff distances of van der Waals and Coulomb forces are set to 1.2 nm in real space under the periodicity assumptions; periodic boundary conditions set in all directions were utilized. All bonds were restrained using the LINCS algorithm. For the production runs, each system was initially simulated for 1 *µ*s, from which two structures selected from the clustering analysis were used as starting structures for the next 2 × 1 *µ*s simulations.

## Acknowledgement

V.F., S.J.L. and H.C. acknowledge VILLUM FONDEN (Villum Young Investigator Grant “Protein Superstructures as Smart Biomaterials, ProSmart”, Grant 19175) and the Novo Nordisk Foundation (NNF20OC0065260 and NNF22OC0080141) for funding the project. We acknowledge the University of Copenhagen, Small angle X-ray facility, CPHSAXS, funded by the Novo Nordisk Foundation (grant no. NNF19OC0055857). https://drug.ku.dk/core-facilities/cphsaxs. The SAXS data using synchrotron radiation were acquired at beamline P12, managed by EMBL Hamburg located at the PETRA III storage ring (DESY, Hamburg, Germany). The authors acknowledge the Core Facility for Integrated Microscopy, Faculty of Health and Medical Sciences, University of Copenhagen. The authors thank the VILLUM FONDEN (Grant 19175) for funding the CLARIOstar plate reader and VILLUM FONDEN (Grant 19175), the Novo Nordisk Foundation (Grant NNF16OC0021948) and the Lundbeck Foundation (Grant R155-2013-14113) for funding the Leica DMi8 microscope. The authors gratefully acknowledge computing time on the supercomputer JURECA at Forschungszentrum Jülich under grant name AMYSALT. M.K. and B.S. are grateful for funding from the Palestinian–German Science Bridge financed by the German Federal Ministry of Education and Research (BMBF).

## Supporting Information Available

Further microscopy, SAXS and MD data are included in the Supporting Information.

---

For Table of Contents Only

**Figure 8:**
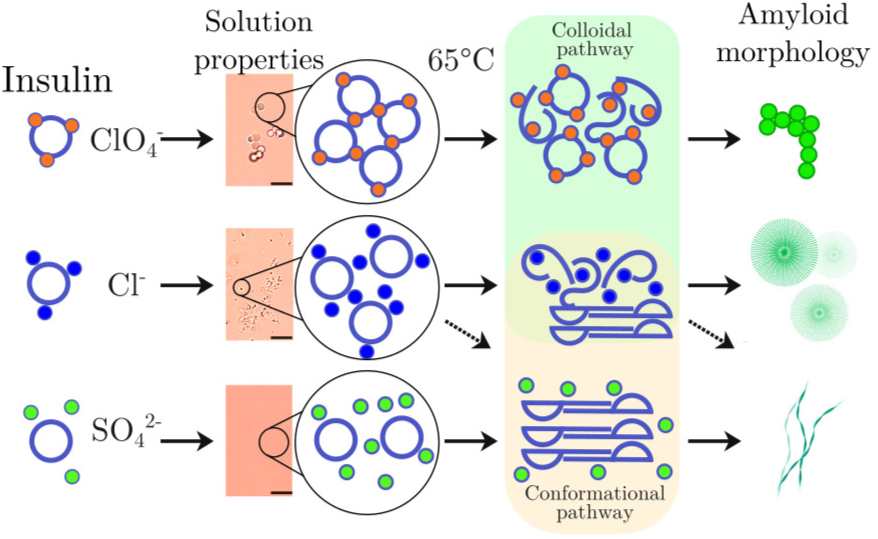
Table of contents.

## Notes

### Competing Interest Statement

The authors have declared no competing interest.

### Summary of Updates

We have included extra data and revised the text

