## Supplementary data for "Insulin Amyloid Morphology is Encoded in H-bonds and Electrostatics Interactions Ruling Protein Phase Separation"

Room Temperature	4°C 30 minutes	Back to room temperature

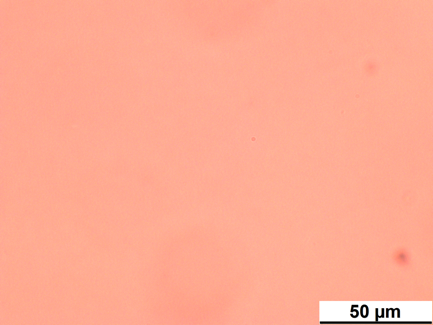

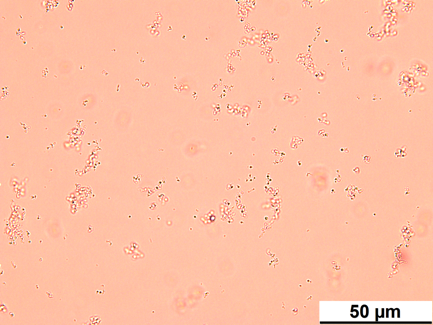

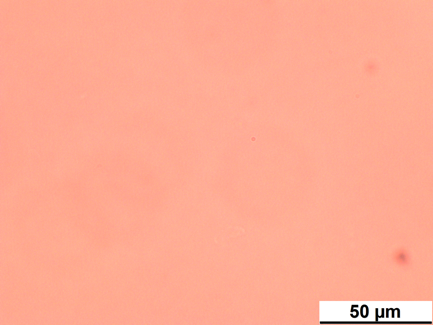

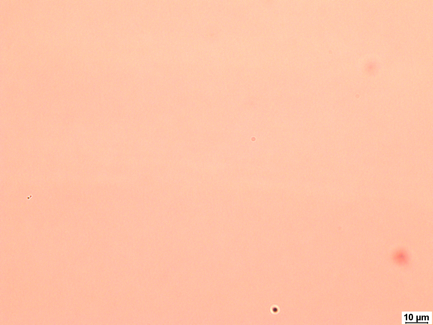

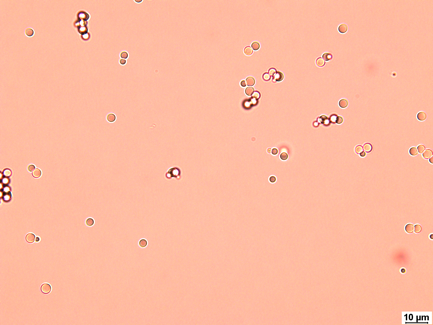

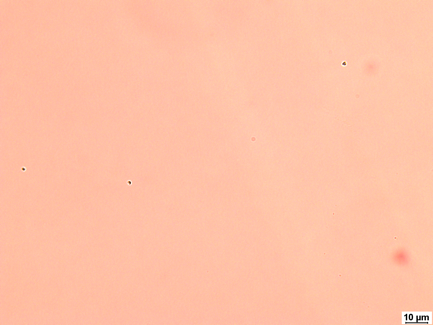

**Figure S1**: Examples of microscope images of the same solutions of 3.4 mg/ml insulin at room temperature, after cooling and heating back to room temperature. Note that for 100 mM NaCl no droplets were observed under any conditions and thus the microscopy images are not shown. Top row: 100 mM ClO_4_, bottom row 100 mM SO_4_^2-^

**Figure S2**: Opacity of insulin (3.4 mg/ml) in 100 mM SO_4_^2-^ at room temperature (Eppendorf on the left) and after incubation at 4°C for 15 minutes (Eppendorf on the right)

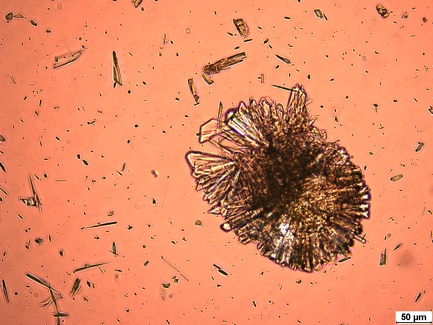

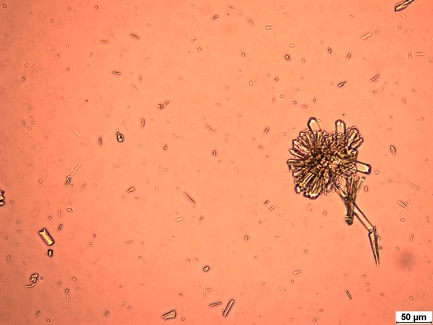

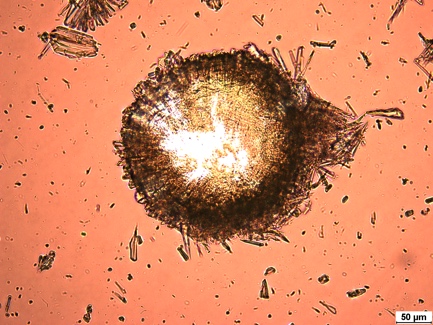

**Figure S3** Protein crystals obtained after incubating 3.4 mg/ml insulin for 48 hours in 100 mM SO_4_^2-^ at room temperature. Images are from different areas of the deposited sample.

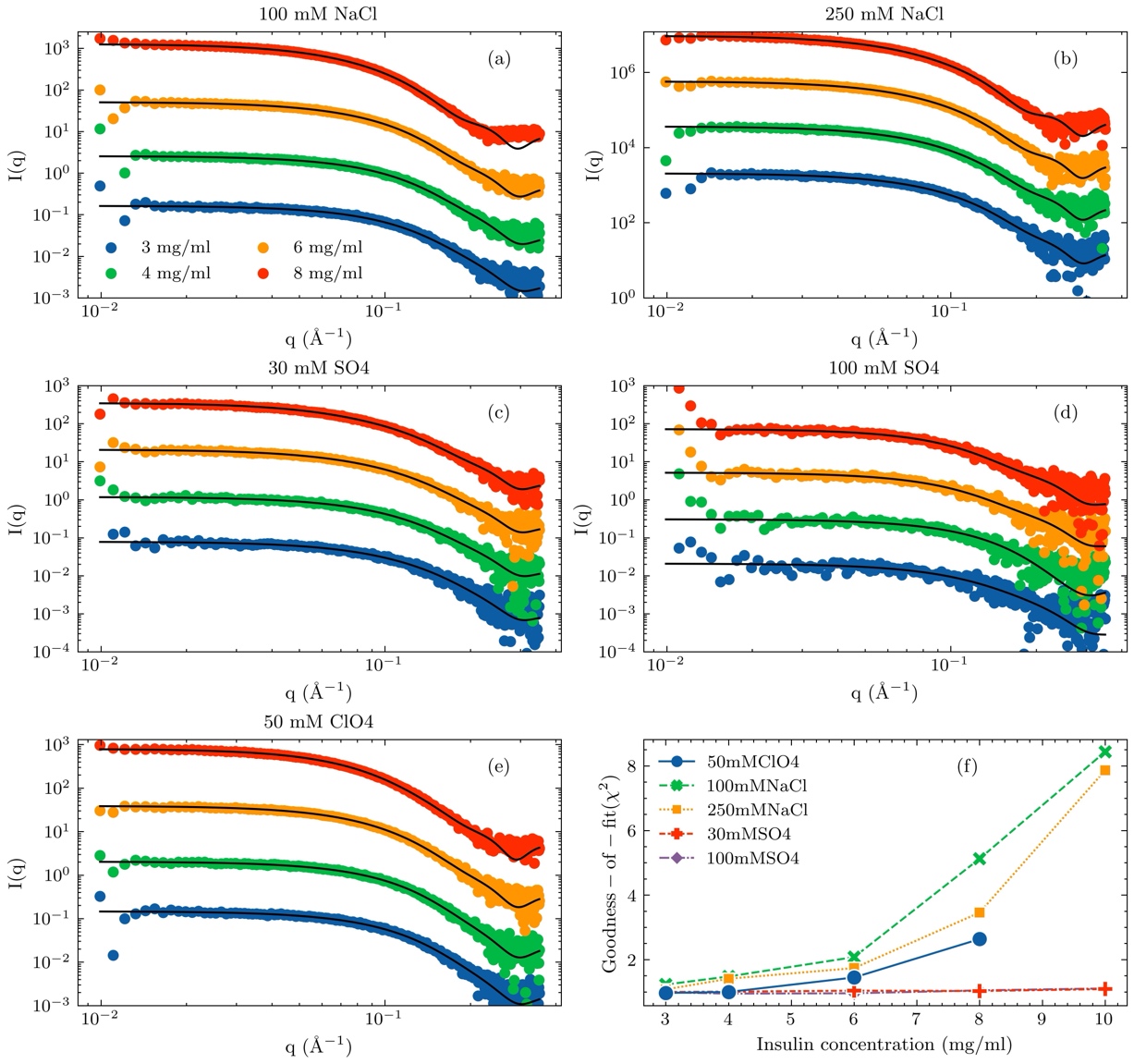

**Figure S4**. (a-e) Fits to the SAXS data obtained using oligomer in the indicated ionic strength conditions, for the indicated anions (f) The goodness of the fit for each anion at the measured insulin concentrations.

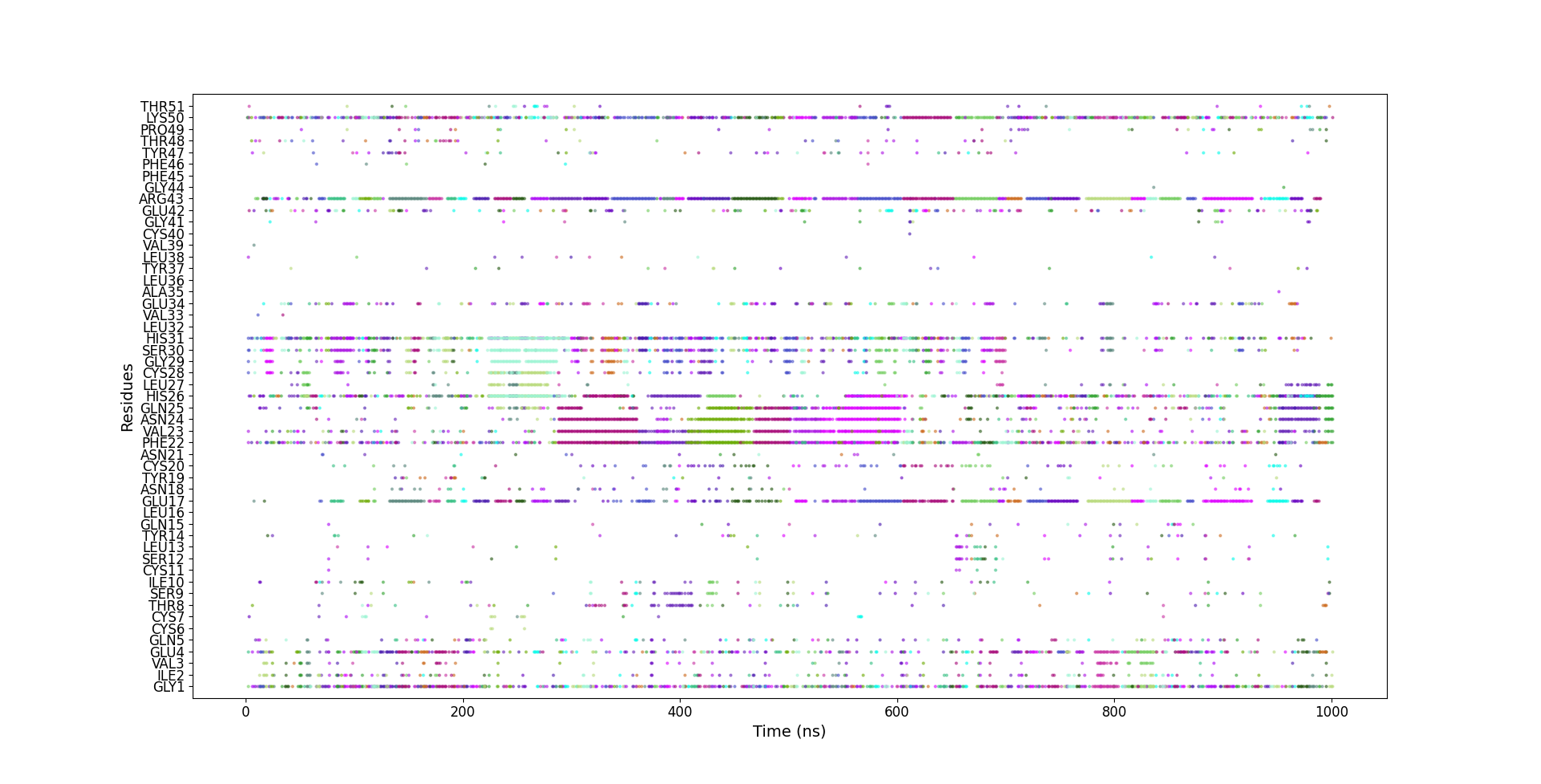
**Figure S5** The ions-protein amino acids contact lifetime of sulphate (cutoff is 0.5 nm) for HI in 100 mM sodium sulphate at 65°C. Each colour represents a different sulphate ion.

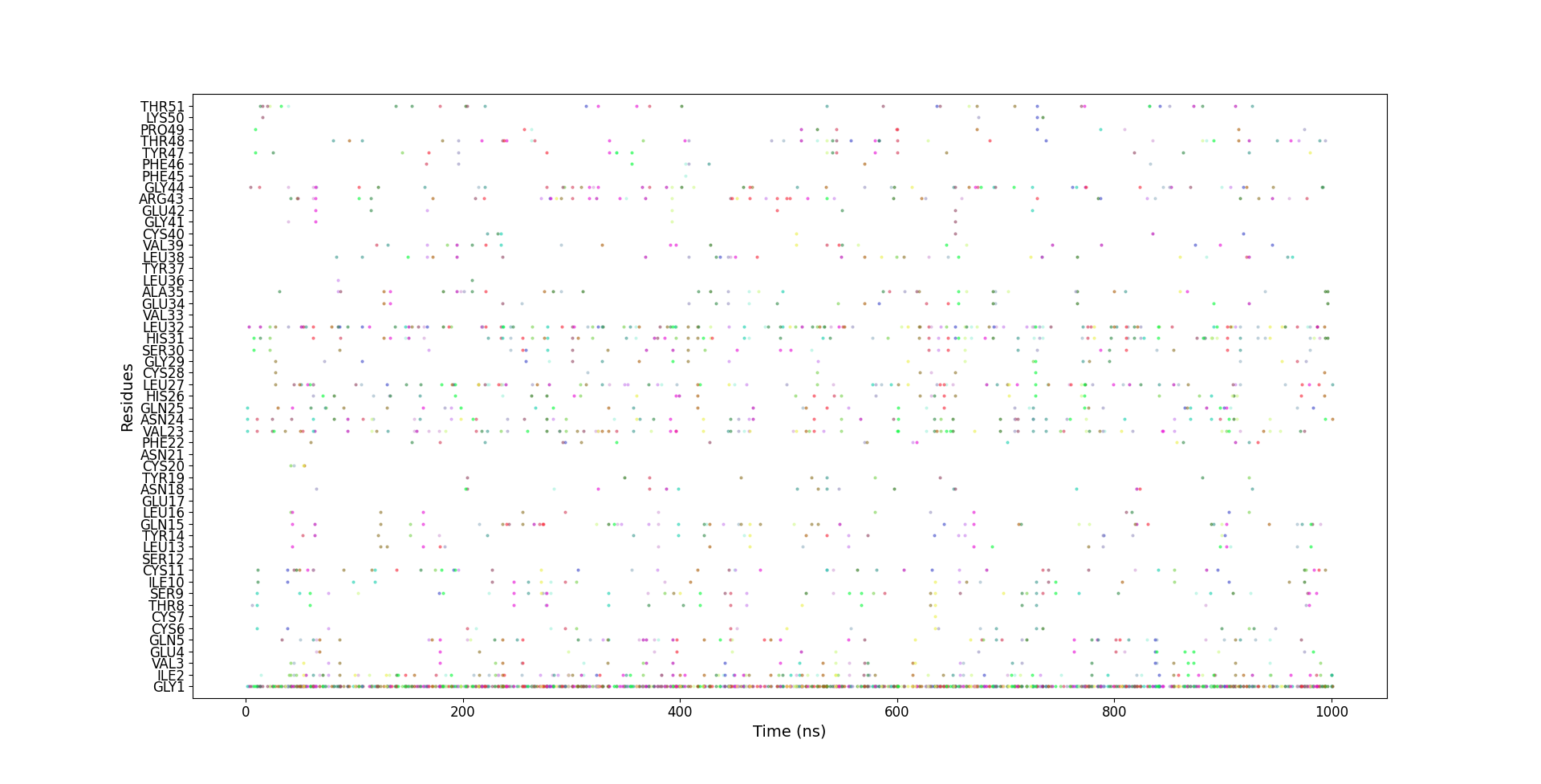

**Figure S6** The ions-protein amino acids contact lifetime of chloride (cutoff is 0.5 nm) for HI in 100 mM sodium chloride at 65°C. Each colour represents a different chloride ion.

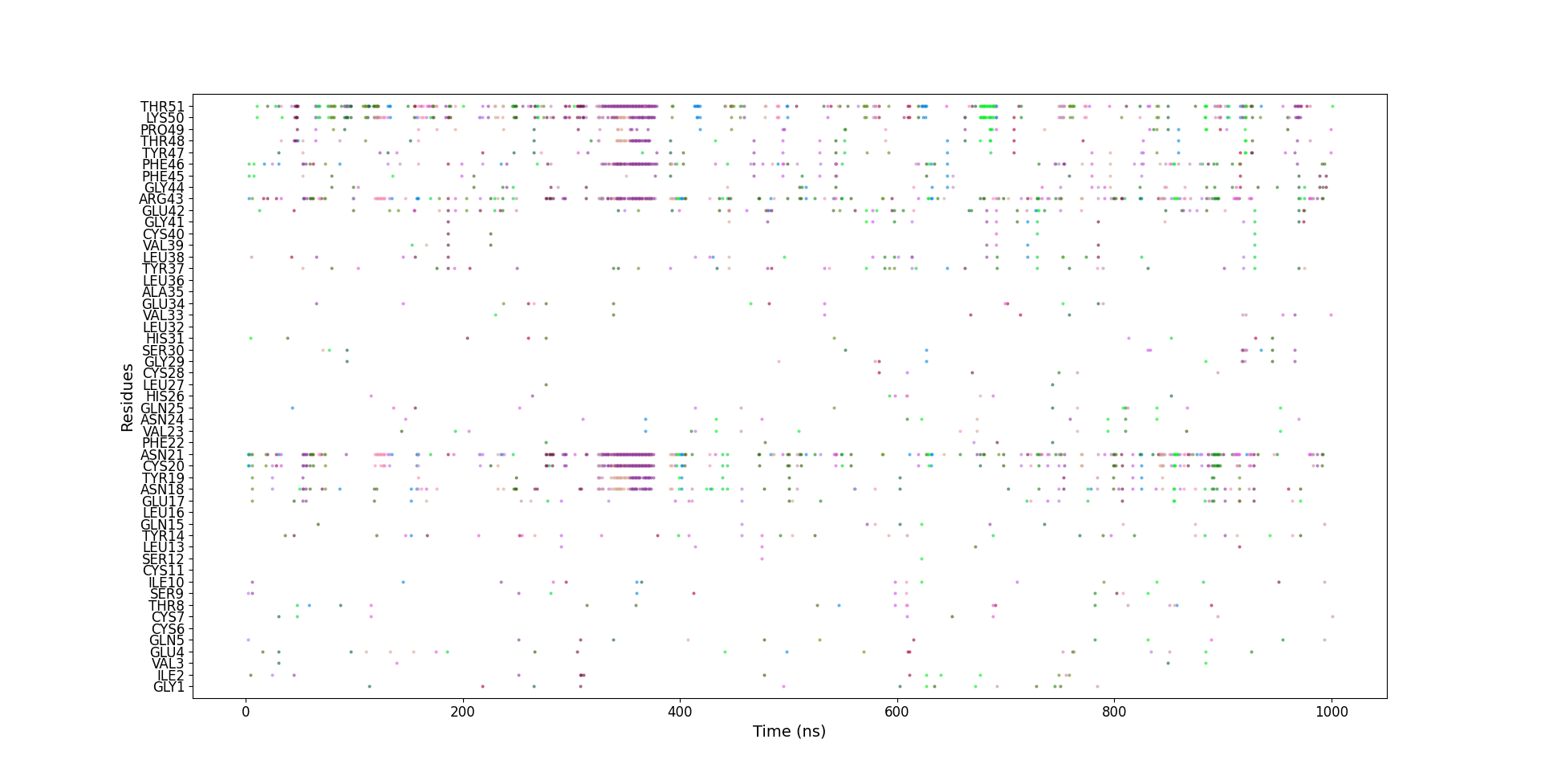

**Figure S7.** The ions-protein amino acids contact lifetime of perchlorate (cutoff is 0.5 nm) for HI in 100 mM sodium perchlorate at 65°C. Each colour represents a different perchlorate ion.
